## Supplementary material for "An open-source, high resolution, automated fluorescence microscope": Rebuttal and additional information

Please find our answers to the reviewers’ comments below in blue:

First, we want to thank the reviewers for their effort to help us improving our work. We apologize for any trivial mistakes we had overlooked. Please find below our answers to the very constructive and helpful comments of the editors.

Reviewer #1 (Recommendations for The Authors):

To complement the current data set:

Figure 2(a & b): Panels i & ii, were chosen on the area where the distribution of the laser appears to be flatter. Can the authors select microtubules from a different section? Otherwise, it is reasonable to also crop the field-of-view along the flatter area (as done in Fig 6).

The figure 2 was changed to according to the reviewer’s suggestions. The profiles of microtubules from a different section have similar profiles, but the region with best illumination thus best SNR of the profile have been used for the figure.

Figure 2(c): The current plot shows the gaussian distribution which does not appear to be centered. Instead of a horizontal line, can the authors provide a diagonal profile across the field of view and update the panel below?

A diagonal cross-section of the illuminated FOV is provided in figure 2 to replace the previous horizontal profile. The pattern seems not to be perfectly radially symmetric, and more light seems to be blocked at the bottom of the illumination pattern compared to the top. A possible improvement can be provided by a fiber-coupled laser, that could provide a more homogeneous illumination while being easier to handle in the assembly process.

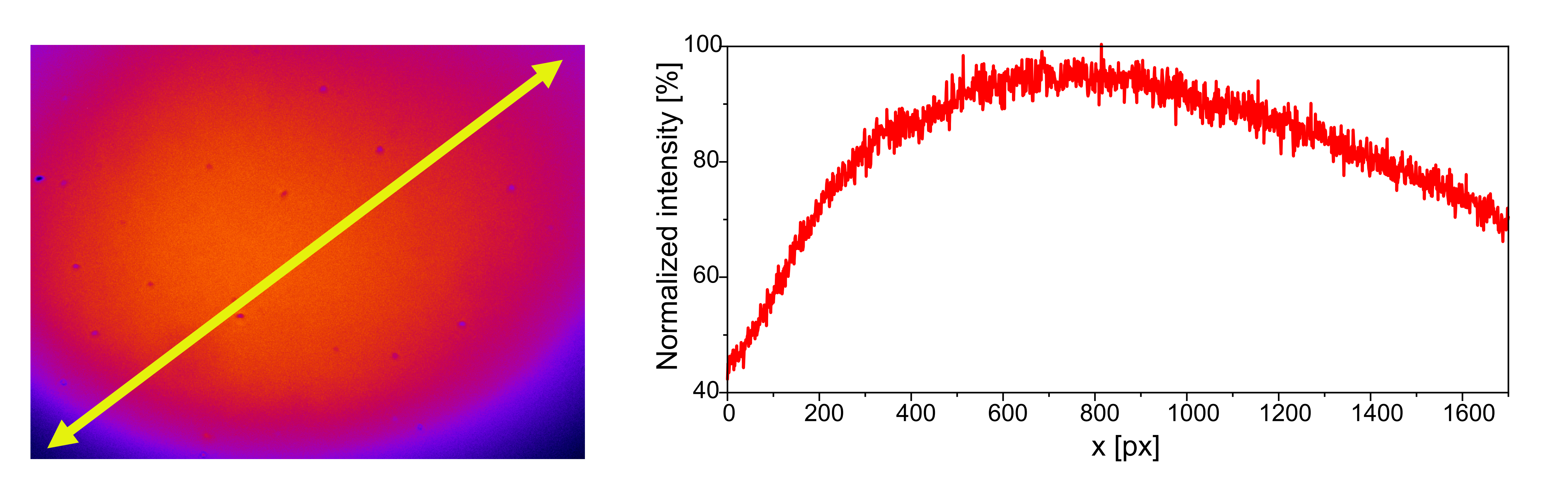

*Diagonal cross-section of the illuminated FOV. Pixel-size (104nm) is the same as in figure 2. Intensity has been normalized according to the maximal value.*

Figure 2(d): The system presents a XY drift of ~500nm over the course of a couple of hours. However, is not clear how the focus is being maintained. Can the authors clarify this point and add the axial drift to the plot?

The axial position of the sample could be maintained over a prolonged period of time without correcting for drift. Measurements where an axial shift was induced by tension pulses in the electronics have been discarded, but the stability of the stage seems to be sufficient to allow for imaging without lateral and axial drift correction. The XY drift measurement displayed in figure 2(d) can be extended by measuring the σ of the PSF over time. The increase of σ would suggest an axial displacement in relation to the focus plane. In these measurements, a slight axial drift can be seen, the fluorescent beads however can still be localized over the whole course of the measurement.

A separate experiment was performed, using the same objective on the UC2 setup and on a high-quality setup equipped with a piezo actuator able to move in 10 nm steps. The precise Z steps of the piezo allows to reproducibly swipe through the PSF shape and to give an estimate of the axial displacement of the sample, according to the changes in PSF FWHM (Full Width at Half Maximum). When superimposing the graph with the UC2 measurement of fluorescent beads with the smallest possible Z step, an estimate about the relative axial position of the sample can be provided. The accuracy of the stage however remains limited.

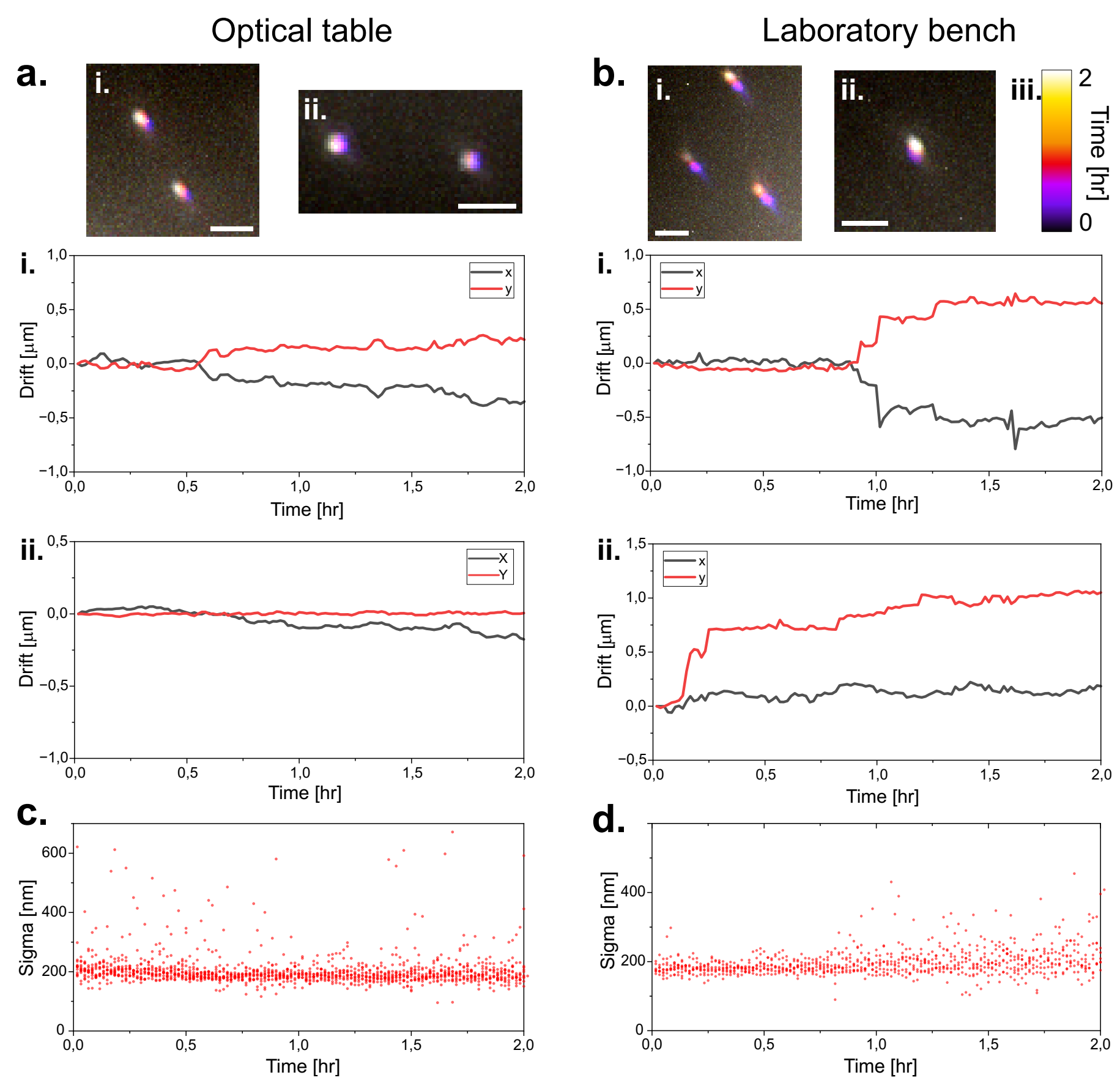

*Drift figure:* *a. Drift of fluorescent TS beads on the UC2 setup positioned upon an optical table over a duration of two hours. Beads are localized and resulting displacement in i. and ii. are plotted in the graphs below. The procedure is repeated in b. with the microscope placed on a laboratory bench instead. c. (for the optical table i.) and d. (for the laboratory bench i.) show the variation in the sigma value of the localized beads over the measurement duration. As the sigma values changes when the beads are out of focus, the stability of the setup can be confirmed, as it remains practically unchanged over the measurement duration.*

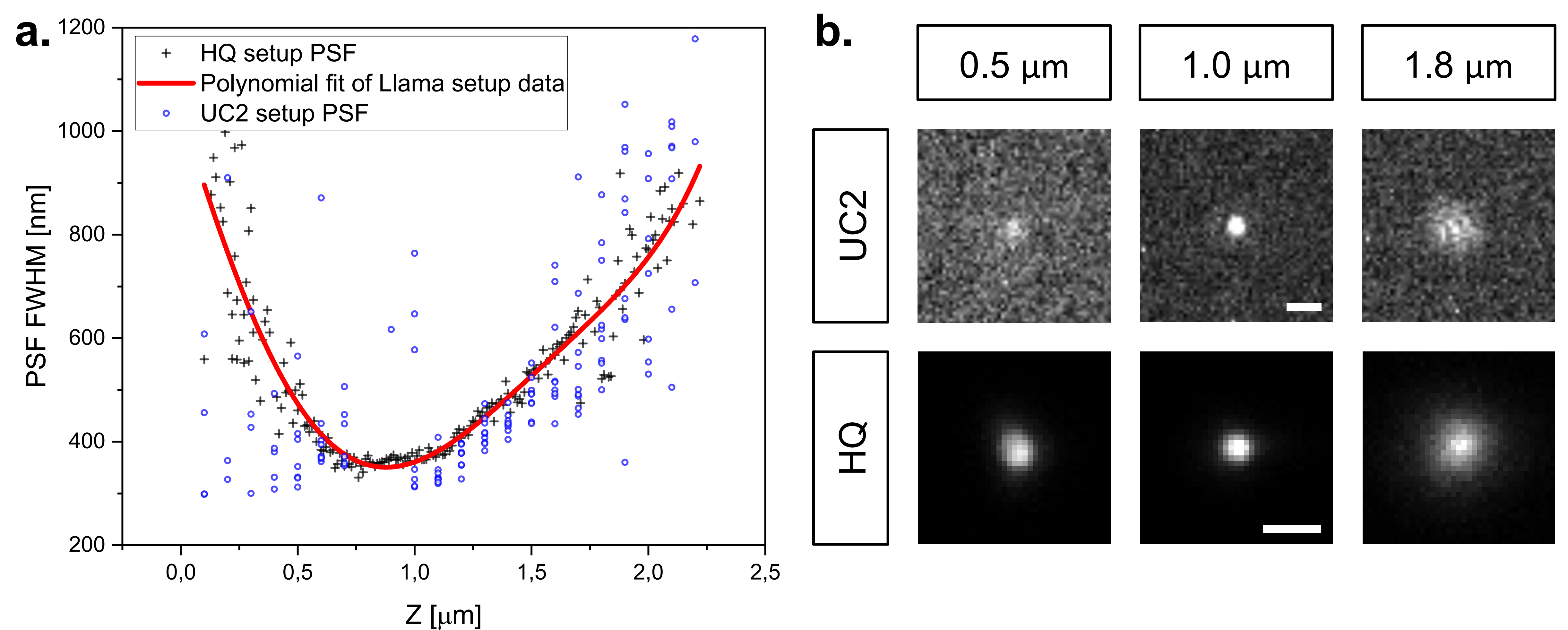

*Z-focus figure: Estimation of the axial position of TS beads on the UC2 setup. a. The change in PSF FWHM was quantified by acquiring a Z stack of a beads sample. The homebuilt high-quality setup (HQ) was used as a reference, by using the same objective and TS sample. The PSF FWHM on the UC2 setup was measured using the lowest possible axial stage displacement. A Z-position can thus be estimated for single molecules, as displayed in b.*

Addressing the seemingly correlated behavior of the X and Y drift:

Further measurement show less correlation between drift in X and in Y. Simultaneous motion in X and Y seems to indicate that the stage or the sample is tilted. The collective movement in X and Y seems accentuated by bigger jumps, probably originating from vibrations (as more predominantly shown in the measurements on the laboratory bench compared to the optical table). Tension fluctuations inducing motion of the stage are possible but are highly unlikely to have induced the drift in the displayed measurements.

Figure 3: Can the authors comment on the effect or otherwise potential effect of the incubator (humidity, condensation etc) may have on the system (e.g., camera, electronics etc)?

When moving the microscope into the incubator, the first precaution is to check if the used electronics are able to perform at 37° C. Then, placing the microscope inside the incubator can induce condensation of water droplets at the cold interfaces, potentially damaging the electronics or reducing imaging quality. This can be prevented by preheating the microscope in e.g. an incubator without humidity, for a few hours before placing it within the functional incubator. The used incubator should also be checked for air streams (to distribute the CO_2_), and a direct exposure of the setup to the air stream should be prevented. The usage of a layer of foam material (e.g. Polyurethane) under the microscope helps to reduce possible effects of incubator vibrations on the microscope. The hydrophilic character of PLA makes its usage within the incubator challenging due to its reduced thermal stability. The temperature also inherently reduces the mechanical stability of 3D printed parts. Using a less hydrophilic and more thermally stable plastic, such as ABS, combined with a higher percentage of infill are the empirical solution to this challenge. Further options and designs to improve the usage of the microscope within the incubator are still in developement.

Figure 5: Can the authors perform single molecule experiments with an alternative tag such

as Alexa647?

The SPT experiments were performed with QDs to make use of their photostability and brightness. The *d*STORM experiment suggests that imaging single AF647 molecules with sufficient SNR is possible. The usage of AF647 for SPT is possible but would reduce the accuracy of the localization and shorten the acquired track-lengths, due to the blinking properties of AF647 when illuminated. The tracking experiment with the QDs thus was a proof of concept that the SPT experiments are possible and allow to reproduce the diffusion coefficients published in common literature. The usage of alternative tags can be an interesting extension of the capabilities that users can perform for their applications.

Figure 6: The authors demonstrate dSTORM of microtubules. It would enhance the paper to also demonstrate 3D imaging (e.g., via cylindrical lens).

The usage of a cylindrical lens for 3D imaging was not performed yet. The implementation would not be difficult, given the high modularity of the setup in general. The calibration of the PSF shape with astigmatism might however be challenging as the vertical scanning of the Z-stage lacks reliability in its current build. Methods such as biplane imaging might also be difficult to implement, as the halved number of photons in each channel leads to losses in the accuracy of localization. As a future improvement of the setup, the option of providing 3D information with single molecule accuracy is definitely desirable and will be tried out. In the following figure, two concepts for introducing 3D imaging capabilities in the detection layer of the microscope are presented.

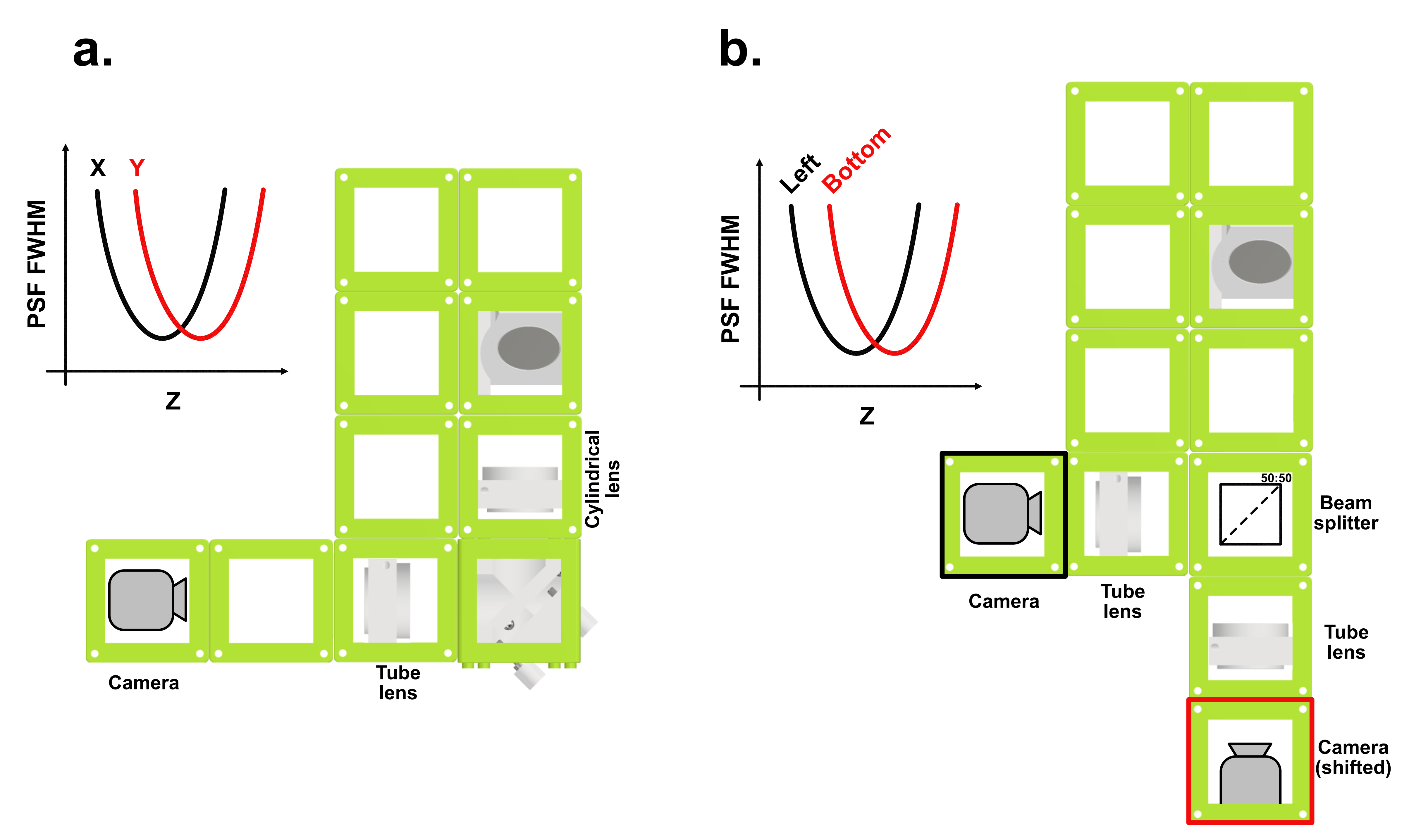

*3D concept figure: Two possible setup modifications to provide axial information when imaging single molecules.* ***a.*** *A cylindrical lens can be placed to induce an asymmetry between the PSF FWHM in x and in y. Every Z position can be identified by two distinct PSF FWHM values in X and Y.* ***b.*** *By splitting the beam in two and defocusing one path, every PSF will have a specific set of values for its FWHM on the two detectors.*

Imaging modalities section: Regarding the use of cling film to diffuse; can the authors comment on the continual use of this approach, including its degradation over time?

The cling foil was only used as a diffuser for broadening the laser profile. A detailed analysis of the constitution of the foil was not done, as no visible changes could be seen on the illumination pattern and the foil itself. The piece of cling foil is attached to a rotor. Detaching of the cling foil or vibrations originating from the rotor need to be minimized. By keeping the rotation speed to a necessary minimum and attaching the cling foil correctly to the rotor, a usable solution can be created. The low price of the cling foil provides the possibility to exchange the foil on a regular basis, allowing to keep the foil under optimal conditions.

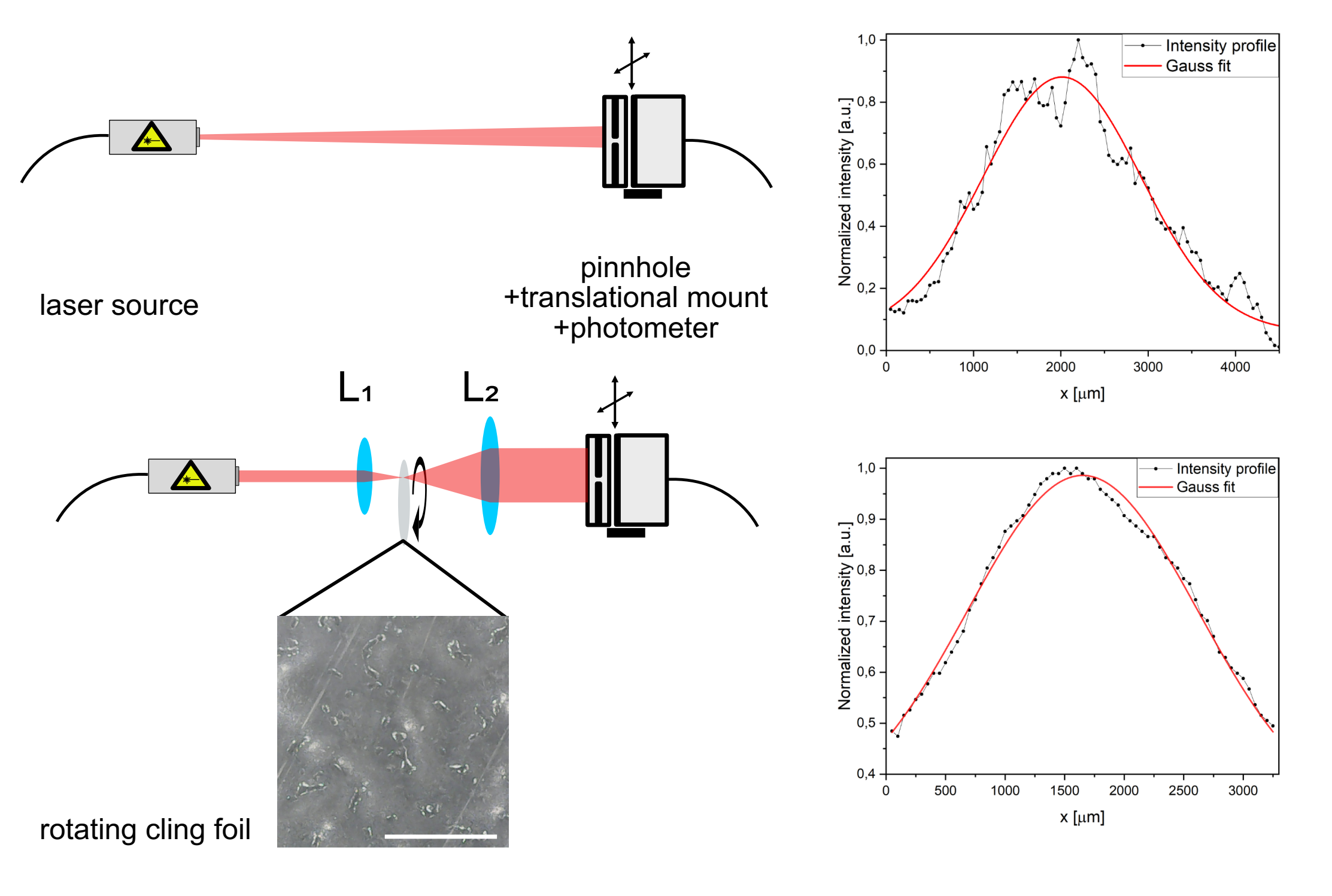

*Profile figure: By moving a combination of pinhole and photometer to scan through the laser profile with a translational mount, the shape of the laser beam can be estimated. The cling foil plays the same role as a diffuser in other setups.*

Reviewer #2 (Recommendations for The Authors):

lines

20, add "," after parts

110, rotating cling foil?

112/116, "custom 3D printed" I thought they were injection molded, please finalize

113, "puzzle pieces" rephrase and they are also barely visible

119, not clear that the stage is a manual stage that was turned into a motorised one by adding belts

123-126, detail for SI,

132, replace Arduino-coded with Arduino-based

143, add reference to Napari

146, (black) cardboard seems to be a cheaper and quicker alternative

153, dichroic

151-155, reads more like a blog post than a paper (maybe add a section on trouble shooting)

156, antibody?

167/189, moderate, please be specific

194, layer of foam material, specify

221, add description/reference to GPI. What is that? why is it relevant?

226: add one sentence description of MMS

318, add "," after students

332-334, as mentioned earlier, not clear, you bought a manual stage and connected belts, correct?

376-377, might be difficult to understand for the layman

391, what laser was used?

figure 1, poor contrast between components, components visible should be named as much as possible, maybe provide the base layer in a different shade. To me, the red and blue labels look like fluorophores.

figure 1. looks like d is the excitation layer and not e, please fix.

figure 2, caption a-c, figure 1-d!, btw, why is the drift so anti-correlated?

figure 6 (line 259) nanometer I guess, not micrometer

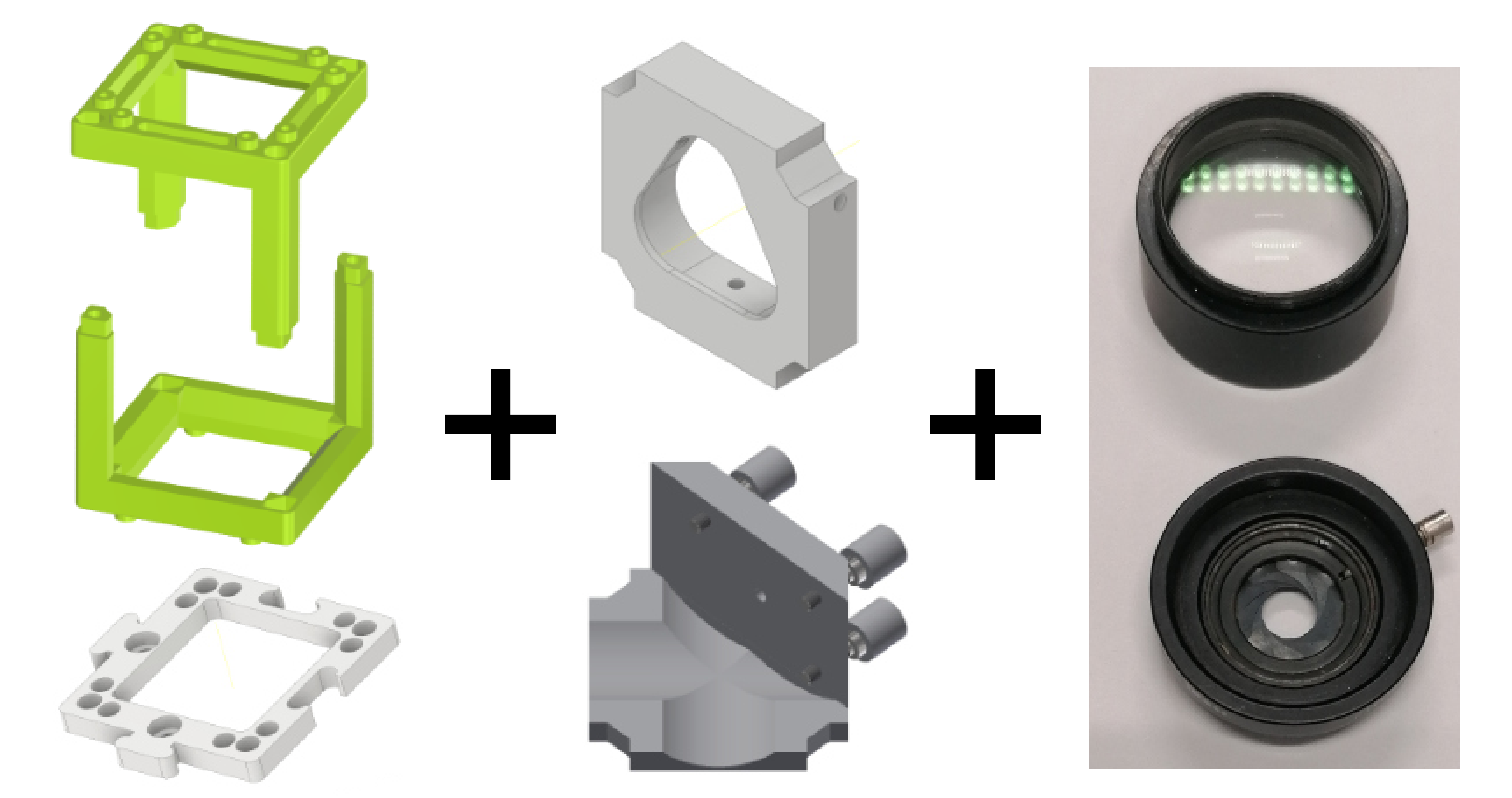

*Figure S1: Basic concept of the UC2 setup: Left: Cubes (green) are connected to one another via puzzle pieces (white). Middle: 3D printed mounts have been designed to adapt various optics (right) to the cube framework. Combined usage of cubes and design of various mounts allows to interface various optics for the assembly.*

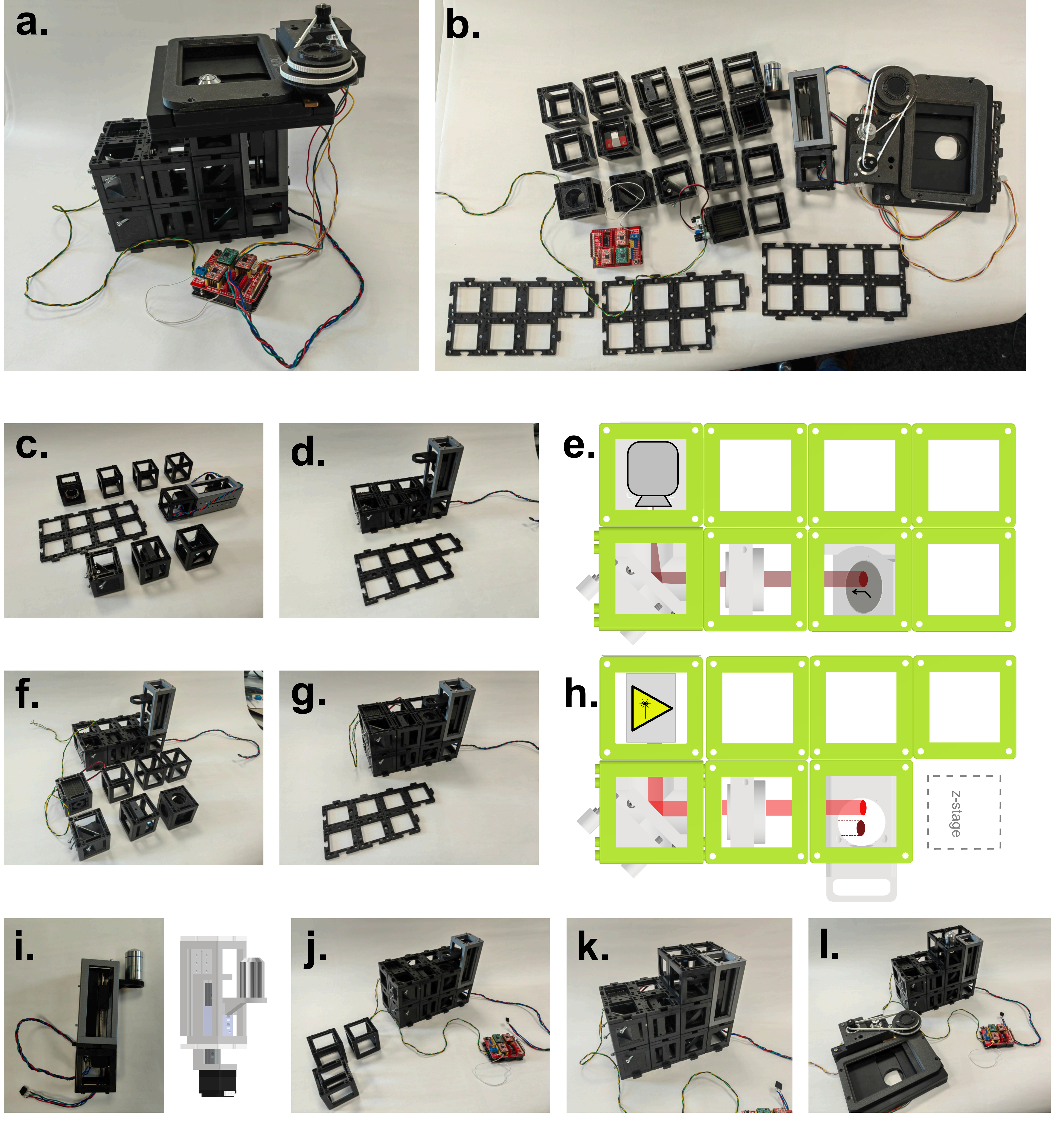

*Figure S2: Building the UC2 widefield microscope:* ***a.*** *Photograph of the complete setup.* ***b.*** *All pieces necessary to build the setup. A list of the components can be found in the bill of materials.* ***c.*** *Bottom emission layer of the microscope before assembly.* ***d.*** *Emission layer after assembly.* *Connection between cubes is doubled by using a layer of puzzles on the top and the bottom of the emission layer.* ***e.*** *CAD schematic of the emission layer and the positioning of the optics.* ***f.*** *Middle excitation layer of the microscope before assembly. Beam magnifier and homogenizer have been left out for clarity.* ***g.*** *Excitation layer after assembly is also covered by a puzzle layer.* ***h.*** *CAD schematic of the excitation layer and the positioning of the optics.* ***i.*** *Z-stage photograph and corresponding CAD file. Motor of the stage is embedded within the bottom cube.* ***j.*** *A layer of empty cubes supports the microscope stage.* ***k.*** *At this stage of the assembly, the objective is screwed into the objective holder.* ***l.*** *Finally, the stage is wired to the electronics and can then be mounted on top of the microscope (see* ***a.****).*

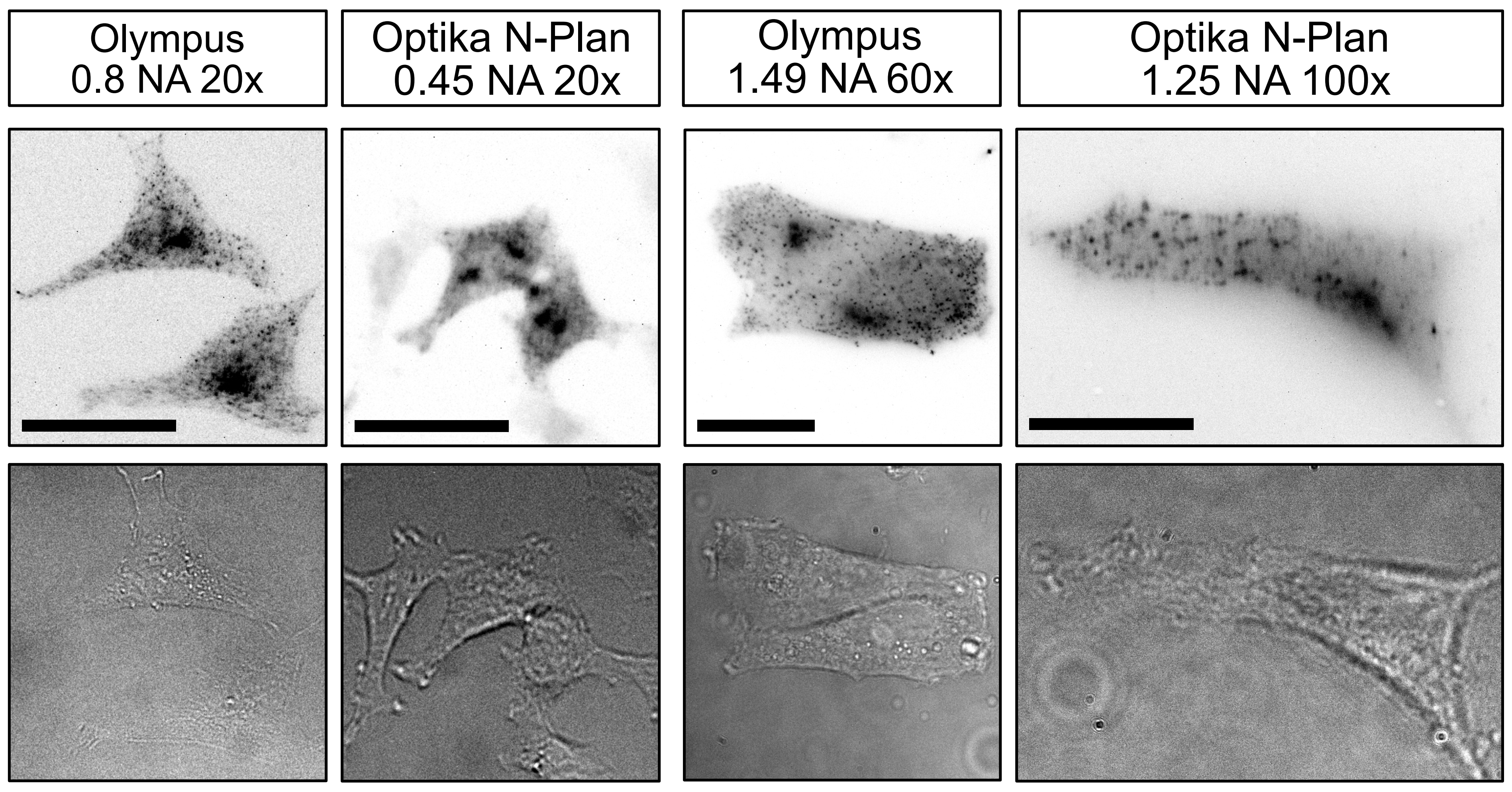

*Figure S3: Measurements performed on the UC2 setup with lower budget objectives. The imaged sample is HeLa cells, stably transfected to express CLC-GFP, then labelled with AF647 through immunostaining. The setup has been kept identical except for the objectives. Scale bar respectively represents 30 µm.*
